# Manta and devil ray genome assemblies uncover signatures of pelagic adaptation

**DOI:** 10.64898/2026.10.27.761883

**Authors:** Léa Soler Clavel, Manu Kumar Gundappa, Atlantine Boggio-Pasqua, Kim Bassos-Hull, Lisa Hoopes, Szu-Hsuan Lee, Marta Gut, Jèssica Gómez Garrido, Fernando Cruz, Tyler S. Alioto, Tatiana Tilley, Nivesh Jain, Jennifer Balacco, Conor Whelan, Dominic E. Absolon, Linelle Abueg, Bonhwang Koo, Giulio Formenti, Olivier Fedrigo, Erich D. Jarvis, Gavin Naylor, Daniel J. Macqueen, Emily Humble

## Abstract

Manta and devil rays (Mobulidae) are iconic marine megafauna highly adapted to life in the open ocean, with forward-facing mouths, enlarged pectoral fins and specialised filter-feeding apparatus. These innovations have enabled mobulids to occupy vast pelagic habitats, yet the genetic changes underlying them remain poorly understood. Here we use a combination of chromatin conformation capture, PacBio HiFi and Nanopore long-reads to generate highly contiguous annotated chromosome-level reference genomes for two mobulid species: the Atlantic pygmy devil ray (*Mobula hypostoma*) and the Atlantic manta ray (*Mobula yarae*). The assemblies span 3.5 and 4.0 Gb, with contig N50s of 23.3 and 7.3 Mb, respectively. We combine these into a Progressive Cactus whole genome alignment with benthic batoid outgroups to characterise the landscape of structural variation and identify candidate molecular regions associated with pelagic adaptation. This approach revealed that over 1.5 million structural rearrangements occurred along the ancestral mobulid lineage, with deletions encompassing three times more sequence compared to the benthic sister lineage. Genes affected by mobulid specific deletions were significantly enriched for neurogenesis pathways linked to craniofacial patterning, with several deletions overlapping the putative promoter sequences of craniofacial and roof of mouth development genes. Our findings highlight candidate developmental pathways through which ancestral deletions may have driven mobulid adaptations, while the high-quality genome resources provide a foundation for comparative and functional studies in Chondrichthyes.

## Introduction

Skates and rays (Batomorphi, Elasmobranchii) are the most diverse group of cartilaginous fishes, with around 600 species described worldwide (Compagno, 2005). While most are benthic, a handful of lineages have adapted to open-water living – a major evolutionary transition associated with changes in locomotion, feeding strategy and ecological niche (Hall et al., 2018). Among the extant pelagic groups, manta and devil rays (Mobulidae, Myliobatiformes) exhibit the most specialised and derived adaptations to life in the open ocean. These innovations have enabled mobulids to successfully colonise pelagic habitats resulting in the most widespread distributions of any ray lineage (Couturier et al., 2012). However, the same traits render mobulids highly susceptible to fisheries exploitation and have contributed to their globally threatened conservation status (Laglbauer et al., 2026). Despite the ecological significance of pelagic specialisation, the underlying genomic and evolutionary processes are poorly understood.

The characteristic morphology of mobulids is integral to their pelagic lifestyle, especially their filter feeding strategy. By unfurling their distinct cephalic fins, mobulids funnel water into their terminal mouths where plankton is filtered through specialised gill plates (Cortés et al., 2008; Sanderson & Wassersug, 1990). Simultaneously, enlarged, wing-shaped pectoral fins provide lift and maneuverability in the water column and support efficient open-water swimming (Fish et al., 2016; Rosenberger, 2001). These morphological traits represent significant deviations from the inferred ancestral batoid condition of a broad, depressed pectoral disk form, likely adapted to a benthic lifestyle (Aschliman et al., 2012).

A handful of studies have explored the genetic basis for mobulid morphological traits (McFarland & Crow, 2024; Swenson et al., 2018), proposing candidate developmental pathways involved in cephalic and pectoral fin formation. However, experimental models to date have relied on either the cownose ray (*Rhinoptera bonasus*, Rhinopteridae) *–* a close relative of mobulids (Swenson et al., 2018) – or gain-of-function assays in little skate (*Leucoraja erinacea,* McFarland & Crow, 2024). Although rhinopterids share certain features with mobulids, such as wing-like fins and cephalic lobes, their craniofacial morphology more closely resembles the likely ancestral batoid body plan, featuring smaller cephalic fins fused anteriorly near a ventrally located mouth (Hall et al., 2018). To gain a more comprehensive understanding of pelagic adaptation, comparative studies of mobulid species themselves are necessary.

Comparative genomic analysis of high-quality, annotated chromosomal-level reference genomes provides a powerful framework to investigate adaptive evolution, underpinned by a rapidly emerging toolbox of whole genome alignment methods (Armstrong et al., 2019). This approach enables detailed exploration of repeat landscapes (Sotero-Caio et al., 2017), chromosomal rearrangements (T. D. Lewin et al., 2024) and structural variation (Jayakodi et al., 2024), all representing established drivers of evolutionary innovation. Recent advances in long-read sequencing have substantially improved genome contiguity and accuracy, facilitating in-depth studies of lineage-specific traits (Christmas et al., 2023; Feng et al., 2020; Shao et al., 2023). Together with global sequencing initiatives (Formenti et al., 2026; H. A. Lewin et al., 2018; Rhie et al., 2021), these advances have accelerated the availability of genomic resources across diverse taxa (Kuderna et al., 2023; T. D. Lewin et al., 2024; Upham & Landis, 2023). Genomics of elasmobranchs has historically lagged behind other vertebrates, leaving gaps in our understanding of their unique biology and evolution (Nishimura et al., 2022). Fortunately, elasmobranch reference genome assemblies are accumulating rapidly, providing, for example, insights into adaptive immunity (Tan et al., 2021; Veríssimo et al., 2023), skeletal development (Venkatesh et al., 2014), fin and limb evolution (Marlétaz et al., 2023), olfaction (Hara et al., 2018; Marra et al., 2019) and sex chromosome evolution (Lee et al., 2025; Niwa et al., 2025).

Like all mobulids, the Atlantic pygmy devil ray (*Mobula hypostoma)* and the Atlantic manta ray (*Mobula yarae*), are fully pelagic, using their pectoral fins for propulsion and their cephalic fins to funnel plankton into their mouths for filter feeding (Figure 1A-B; Figure S1, Boggio-Pasqua et al., 2026; Couturier et al., 2012). These two species are among the most coastal and reliably encountered mobulids, particularly around Florida, where annual tagging and observational studies have yielded valuable insights into their population dynamics and ecology (Boggio-Pasqua et al., 2026; Bucair et al., 2025; Humble et al., 2025; Pate & Marshall, 2020). Both species are also maintained in *ex situ* collections in North America. This level of accessibility offers a rare opportunity for repeated, high-quality sampling to enable comparative genomic analysis. In this study, we combined Nanopore long-read, PacBio HiFi and Hi-C chromatin conformation capture sequencing to produce highly contiguous chromosomal-level reference genomes for both species, which were annotated using multi-tissue RNA-seq data. We performed a multi-species whole genome alignment with other available batoid genomes to characterise synteny and structural rearrangements across the group. Finally, we leverage these resources to explore lineage-specific deletions as a potential substrate for phenotypic evolution during the transition of mobulids to pelagic environments.

**Figure 1.**
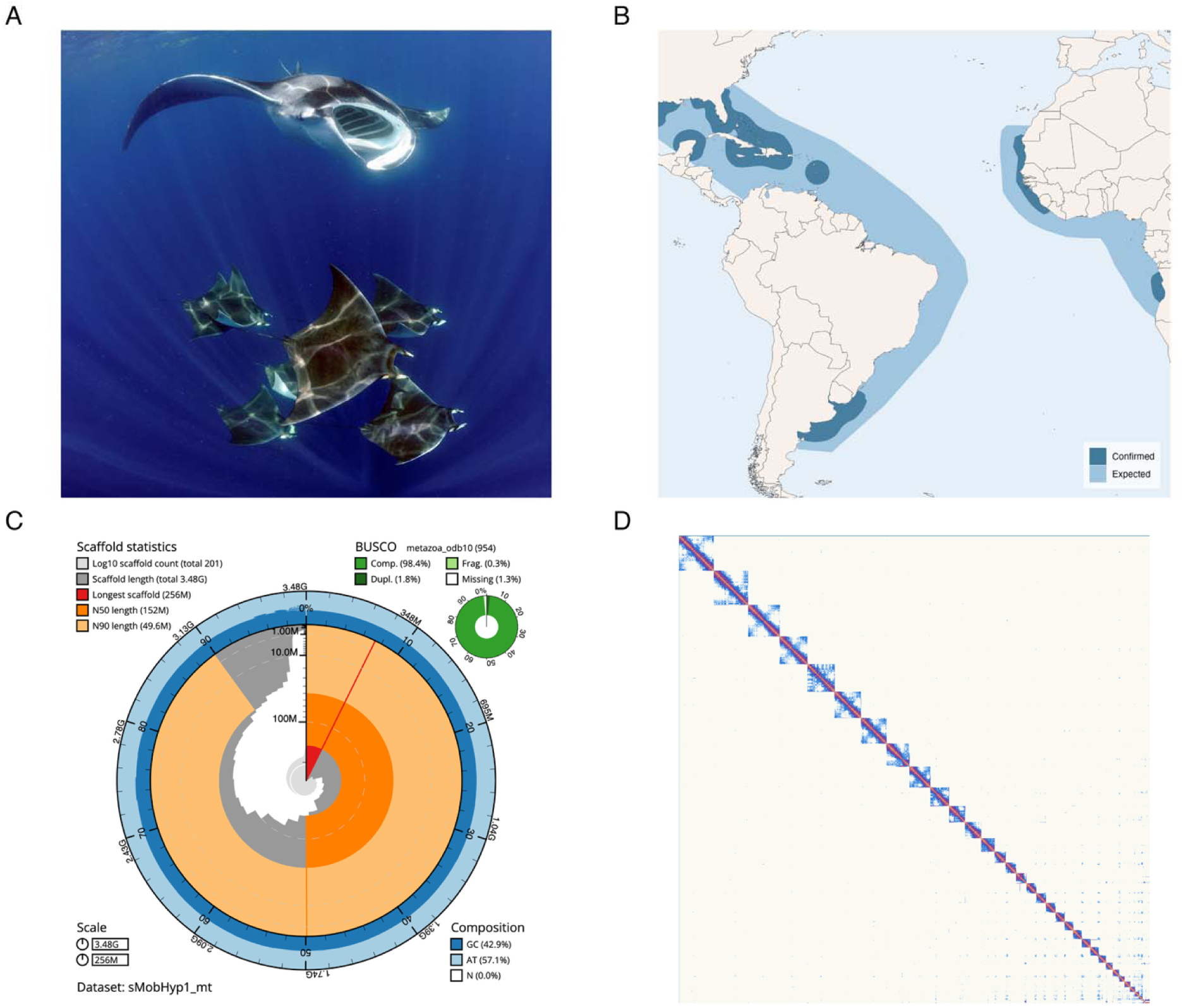
(A) Atlantic pygmy devil rays (*Mobula hypostoma*) and an Atlantic manta ray (*M. yarae*), photographed in Isla Mujeres, Mexico, copyright Guy Stevens. (B) Geographic distribution of *M. hypostoma*. Dark-shaded areas denote the confirmed species range, and light-shaded areas denote the expected species range. See Figure S1 for the geographic distribution of *M. yarae*. (C) Snail plot generated using BlobToolKit to visualize key quality metrics of the *M. hypostoma* genome assembly. Tracks include scaffold count, scaffold length, scaffold N50/N90, GC content and BUSCO scores. (D) Hi-C contact map for the assembled chromosomes in the *M. hypostoma* genome assembly. Chromosomes are ordered and orientated along the axes and the heatmap shows contact frequencies with the strong diagonal signal indicating high contiguity and correct orientation of chromosome-length scaffolds.

## Materials and Methods

### Sample collection

Whole blood for *M. hypostoma* genome sequencing was collected from a juvenile male in Navarre, Florida, USA, in November 2022. The individual was captured using a nylon seine net and transferred to a research vessel, where blood was drawn from the pectoral fin using a sterile syringe. The blood sample was stored in a 2 mL EDTA-K2 tube and immediately placed in liquid nitrogen. All live animal handling procedures were conducted under Florida Protected Species permits SAL-22-1140A-SRP and were approved by the Mote Marine Laboratory Institutional Animal Care and Use Committee (IACUC approval 22-09-KBH2). The sample was transferred between Convention on International Trade in Endangered Species of Wild Fauna and Flora (CITES) registered scientific institutions (19US20116D/9 and ES006) and forwarded to Centro Nacional de Análisis Genómico (CNAG) in Barcelona, Spain for sequencing. For *M. hypostoma* RNA sequencing, samples from heart, ovary, spleen, kidney, liver, muscle, brain, intestine and gill were collected from a deceased adult female housed at Georgia Aquarium, Atlanta, USA. Samples were placed into cryovials within one hour post-mortem and immediately frozen at -80°C before shipment to Novogene, USA, for RNA sequencing.

For manta ray genome and RNA sequencing, samples were collected from a recently deceased adult female housed at Georgia Aquarium and originally caught off southwestern Florida. At the time of sample collection and sequencing, this individual was suspected to represent a putative third species of manta ray (*M. cf. birostris*). This species has since been formally described as the Atlantic manta ray (*Mobula yarae*), using morphological data from the sampled individual (Bucair et al., 2025), and we therefore refer to it here as such. However, we note that species boundaries within the manta ray complex would benefit from further investigation. Samples from heart, spleen, ovary, muscle, liver and kidney were placed into cryovials within four hours post-mortem and immediately frozen at -80°C before shipment to the Vertebrate Genomes Lab (VGL) at the Rockefeller University, USA for genome and RNA sequencing. For all Georgia Aquarium specimens, neither research permits nor CITES documentation were required as samples were collected post-mortem and shipped domestically.

### Generation of Mobula hypostoma sequencing data

High-molecular weight genomic DNA was extracted from fresh-frozen *M. hypostoma* blood using the Qiagen Genomic-tip 20/G kit following the manufacturer’s protocol. DNA quantity and purity were assessed using a Qubit DNA BR Assay Kit (Thermo Fisher Scientific, USA) and Nanodrop2000. DNA integrity was evaluated using the Agilent Femto Pulse system. Paired-end libraries for Illumina sequencing were prepared using a PCR-free protocol with the KAPA HyperPrep Kit (Roche, Switzerland). Genomic DNA underwent end-repair and adenylation, followed by ligation of dual index adaptors (Integrated DNA Technologies, USA). Library quality was assessed on an Agilent 2100 Bioanalyzer and quantified using the KAPA Library Quantification Kit. Sequencing was performed on an Illumina NovaSeq 6000 platform (2 x 151 bp), yielding 134 Gb of raw data.

Oxford Nanopore Technologies (ONT) libraries were prepared using the SQK-LSK114 sequencing kit. Genomic DNA was repaired and adenylated using the NEBNext FFPE DNA Repair Mix (New England Biolabs, NEB, USA) and the NEBNext Ultra II End Repair/dA-Tailing Module (NEB), followed by adaptor ligation. Six sequencing runs were performed on a PromethION 24 using FLO-PRO114M flow cells, with sequencing runtime set to 90 hours per run. Sequencing performance was monitored using MinKNOW v22.12.5 and basecalling was performed using Guppy v6.4.6 (Wick et al., 2019). Sequencing yielded 23,674,327 reads (332 Gb of basecalled data) with N50 of 24 kb length, and a mean length of 14 kb (Figure S2A). Approximately 50 Gb of data (15%) were contained in reads longer than 50 kb. Chromatin conformation capture libraries were generated using the Dovetail Omni-C Kit with the frozen blood samples as input. Chromatin was cross-linked with formaldehyde (Sigma-Aldrich, USA), digested with DNase I, and proximity-ligated following end-repair and ligation of biotinylated bridge adapters. After reverse crosslinking, DNA was purified and Illumina sequencing libraries prepared without fragmentation. Biotinylated fragments were enriched using streptavidin beads and the final library was PCR-amplified using KAPA HiFi HotStart ReadyMix (Roche). The library was sequenced on an Illumina NovaSeq 6000 (2 x 150 bp, dual indexing), generating 185 Gb of raw data. For RNA sequencing, poly(A)-enriched RNA libraries were prepared from the nine *M. hypostoma* organ samples using the NEBNext® Ultra™ II RNA Library Prep Kit (Novogene, USA). Libraries were sequenced on an Illumina NovaSeq 6000 platform, yielding 112.7 Gb of raw data, with 60–193 million 150 bp paired-end reads per sample. Quality control and filtering were performed by Novogene, resulting in >98% of reads retained as clean data.

### Generation of Mobula yarae sequencing data

High molecular weight genomic DNA was extracted from a 13 mg aliquot of frozen *M. yarae* heart tissue. First, the tissue was disrupted and homogenized with a Qiagen TissueRuptor II (Qiagen, Netherlands; Cat. No. 9002755), and then DNA was isolated and cleaned using a Circulomics Nanobind Tissue Big DNA Kit (Circulomics, USA; NB-900-701-01). Bionano optical mapping was performed by labelling 750 ng of DNA with direct labeling enzyme (DLE1) and the Bionano Prep Direct Label and Stain (DLS) protocol (Document number 30206). The labelled DNA was imaged using a Bionano Saphyr. Separately, 10 µg DNA was fragmented to 15 kb – 20 kb using a Megaruptor 3 instrument (Diagenode, USA). A PacBio SMRTbell library was prepared with 5 µg of sheared DNA and SMRTbell Express Template Prep Kit 2.0 (PacBio, USA; PN 100-938-900). The SMRTbell library was size-selected with a Pippin HT instrument (Sage Science, USA) to remove fragments under 10 kb. The size-selected library was then assessed for quantity and quality using a Qubit 3 Fluorometer (Invitrogen, USA; Cat. No. Q33216) and an Agilent Fragment Analyzer. The library was then sequenced on a PacBio Sequel IIe instrument with 30-hour movie time, Sequel Binding Kit 2.2 (PacBio; PN:101-894-200), and Sequel II 8M cells (PacBio; PN: 101-389-001), yielding 113 Gb of raw data. A subsection of frozen heart tissue was shipped to the Arima Genomics sequencing facility (Arima Genomics, USA) to generate an Arima Hi-C library using the Hi-C kit 2.0 and then sequenced on the Novaseq 6000 platform, yielding 584 Gb of raw data. To generate transcriptomic data for annotation, total RNA was isolated from frozen liver, heart, muscle, spleen, ovary, and kidney using a Qiagen RNeasy kit (cat. 74104). RNA integrity score (RIN) was assessed using an Agilent Fragment Analyzer and an RNA-seq library was prepared for each tissue following the Illumina Stranded mRNA Prep kit. The RNA-seq libraries were sequenced on an Illumina NovaSeq 6000, yielding 75 Gb of raw data, with 193–308 million 100 bp paired-end reads per sample.

### Reference genome assemblies and annotation

The chromosome-level genome assembly for *M. hypostoma* was generated using the CLAWS pipeline v2.1 (Figure S3, Gomez-Garrido, 2024), integrating ONT long-read sequencing, Illumina short-read sequencing for base accuracy improvement and Omni-C proximity ligation data for chromosomal scaffolding. Prior to assembly, adaptors were removed from the Illumina data using TrimGalore v0.6.7 (https://github.com/FelixKrueger/TrimGalore) with cutadapt v4.1 parameters -j 4 -e 0.1 -q 20 -O 1 -a AGATCGGAAGAGC, and a k-mer database (k=21) was constructed using Meryl v1.3 (https://github.com/marbl/meryl) to estimate genome characteristics (haploid genome size, heterozygosity and repeat content) with Genomescope2 (Ranallo-Benavidez et al., 2020). ONT reads were quality filtered using Filtlong v0.2.1 (--minlen 1000 --min_mean_q 80 -t 300000000000, https://github.com/rrwick/Filtlong, Figure S2B) and assembled using Nextdenovo v2.5.0 (Hu et al., 2024) in nano-raw mode with an estimated genome size of 5 Gb and otherwise default parameters. The assembly was then polished with HyPo v1.0.3 (Kundu et al., 2019), using both Illumina reads (-b) and ONT reads (-B), with the coverage for Illumina reads set to 75 (-c 75.45164225490777), 100 threads (-t 100) and a batch size of 6 (-p 6). Alternative haplotypes and redundant repetitive regions were removed from the polished assembly using purge_dups v1.2.6 (Guan et al., 2020, Table S1), collapsing the assembly into a single haploid (primary) representation. The assembly was scaffolded with Omni-C data using YaHS v1.2a (Zhou et al., 2023) and manually curated in PretextView v0.2.5 (https://github.com/wtsi-hpag/PretextView), resulting in one contig cut, four gap breaks, and sixteen joins. Assembly consensus accuracy quality value (QV) was assessed using Merqury v1.3 (Rhie et al., 2020) re-using the Illumina WGS read 21-mer meryl database constructed previously. Potential contamination was identified using BlobToolKit v3.1.0 (Challis et al., 2020) running BUSCO (Benchmarking Universal Single-Copy Orthologs) v5.3.2 (Simão et al., 2015) by comparison against the NCBI nt (updated May 26, 2023), UNIPROT reference proteomes (Release 2023_02, 03-May-2023), and multiple BUSCO odb10 databases (*Vertebrata, Metazoa, Eukaryota, Fungi*, Kriventseva et al., 2019).

The chromosome-level genome assembly for *M. yarae* was generated by integrating the HiFi and Hi-C data using the VGP Pipeline 2.0 described in (Larivière et al., 2024). Briefly, the assembly used HiFiasm-HiC mode (0.16.1+galaxy2) resulting in two complete haplotypes (sMobBir1.hap1 and sMobBir1.hap2). Scaffolding with Hi-C data was performed with YaHS (1.2a+galaxy1). To manually curate the assemblies, Hi-C contact maps were generated and manipulated in *PretextView* to correct assembly structural errors and identify chromosomes.

Sex chromosomes were identified using sequencing coverage for both species. The presence of a second X chromosome in *M. yarae* was determined based on comparative alignment to *M. hypostoma* (see Results and Discussion). Autosomes were named in order of decreasing size for both species. Telomere identification was carried out on all assemblies using an in-house script that scans the terminal 100 kb of each scaffold ≥ 5 Mb for the canonical vertebrate telomeric motif (TTAGGG/CCCTAA). A scaffold end was classified as telomeric if repeat density exceeded a threshold of 10 per 1,000 bp in any window within the 100 kb terminal region. Contiguity metrics were generated for all assemblies using gfastats v1.3.12 (Formenti et al., 2022). The primary *M. hypostoma a*ssembly (sMobHyp1.1) and the *M. yarae* haplotype 1 assembly (sMobBir1.hap1) were annotated using the NCBI RefSeq Eukaryotic Genome Annotation Pipeline which used evidence from the RNAseq data produced as part of this project (*M. hypostoma* BioProject: PRJEB71148; *M. yarae*: BioProject: PRJNA1218823) together with protein alignments from a suite of related species (via ProSplign) ton predict the final gene models using Gnomon.

### Mitogenome assembly

The *M. hypostoma* mitochondrial genome was assembled using the FOAM v0.3 pipeline (https://github.com/cnag-aat/FOAM), leveraging a reference-guided approach with ONT and Illumina sequencing data. Mitochondrial ONT reads were selectively captured using the *M. mobular* mitogenome (NC_018784.1, Poortvliet et al., 2015) as a reference and assembled into a circular contig with Flye v2.9 (Kolmogorov et al., 2019). The resulting contig underwent two rounds of Illumina-based polishing using NextPolish v1.4.1 (Hu et al., 2020). Annotation was conducted using MITOS v2.1.3 (Bernt et al., 2013) and the final sequence was oriented and rotated to start at the trnF gene. Mitochondrial genome assembly was attempted for *M. yarae* using MitoHifi v2.2 (Uliano-Silva et al., 2023). Candidate mitogenome sequences were recovered from the HiFi reads however did not align with expected reference sequences and could not be confidently annotated with MITOS. This likely reflects depletion of high-quality mitochondrial sequences in the HiFi library due to high molecular weight DNA fragmentation and size-selection (Formenti et al., 2021). For this reason, no mitogenome is reported for *M. yarae*.

### Progressive Cactus whole genome alignment

To form a backbone for our subsequent analysis, we constructed a reference free multi-species whole genome alignment using the newly assembled *M. hypostoma* and *M. yarae* (sMobBir1.hap2) genomes and publicly available assemblies from three other batoids with annotated chromosome-level genome assemblies available on NCBI: Atlantic stingray *Hypanus sabinus* (Accession number: GCA_030144855.1), thorny skate *Amblyraja radiata* (Accession number: GCA_010909765.2) and little skate *Leucoraja erinacea* (Accession number: GCA_028641065.1, (Marlétaz et al., 2023). These species were chosen to represent benthic myliobatiform and rajiform lineages and serve as outgroups for comparative genomic analysis (Table 1; Figure 3A). Whole-genome alignment was performed using Progressive Cactus v2.7.2 (Armstrong et al., 2020), which also reconstructs ancestral genomes at nodes in a pre-defined phylogenetic tree. The phylogeny used to guide the alignment (Figure 3A) reflected published batoid phylogenies and divergence estimates (Aschliman et al., 2012; Valsecchi et al., 2005; Villalobos-Segura & Underwood, 2020). Four ancestral nodes were specified: Anc0 (ancestral genome for all species); Anc1 (common ancestor of little skate and thorny skate); Anc2 (common ancestor of Atlantic stingray and mobulids); and Anc3 (common ancestor of mobulids) (Figure 3A).

**Table 1.** Quality metrics generated using gfastats v1.3.12 for the five focal batoid assemblies used in this study.

| Genome assembly | Assembly name | Length (Gb) | Assembled chromosomes | Number of scaffolds | Scaffold N50 (Mb) | Number of contigs | Contig N50 (Mb) | Gap length (bp) |
| --- | --- | --- | --- | --- | --- | --- | --- | --- |
| *M. hypostoma* | sMobHyp1.1 | 3.48 | 34 | 200 | 152.4 | 897 | 23.3 | 139,400 |
| *M. yarae*<br>*haplotype 1* | sMobBir1.hap1 | 4.01 | 33 | 4,715 | 148.7 | 6,525 | 7.3 | 30,852,630 |
| *M. yarae*<br>*haplotype 2* | sMobBir1.hap2 | 3.63 | 33 | 1,558 | 152.4 | 3,300 | 8.6 | 54,983,815 |
| *H. sabinus* | sHypSab1.hap1 | 4.04 | 35 | 2,989 | 112.3 | 4,637 | 4.3 | 329,600 |
| *A. radiata* | sAmbRad1.1.pri | 2.56 | 49 | 957 | 62.1 | 4,710 | 1.4 | 190,972,533 |
| *L. erinacea* | Leri_hhj_1 | 2.19 | 50 | 1,769 | 57.2 | 2,631 | 5.2 | 81,010 |

### Comparative analyses of repeat and synteny across myliobatiforms

Repeat content was compared using a standardised approach across the five batoid species to assess contributions to genome size and to compare major repeat families among species. RepeatModeler v.2.0.5 and RepeatMasker v.4.1.7 (Smit et al., 2013) were used in each case to create a *de novo* repeat library per species and produce a repeat-masked version of the *M. hypostoma* genome assembly. Repeats were classified into DNA transposons, retroelements, tandem repeats and unclassified repeats. To determine the extent of chromosome conservation and reorganisation among batoids, we characterised synteny using our Progressive Cactus alignment. Syntenic regions were extracted from the Cactus alignment HAL file using the halSynteny command from HalTools v2.6.12 (Hickey et al., 2013) for comparisons between *M. hypostoma* and *M. yarae*, *M. hypostoma* and *L. erinacea* and between *M. yarae* and *H. sabinus* genomes. *Amblyraja radiata* was not included in this analysis. Synteny blocks were restricted to chromosome-level scaffolds only. Syntenic relationships were visualised as a ribbon plot using the graphics.karyotype module of jcvi v1.6.7 (Tang et al., 2024). Species were arranged in order, in order to keep the two mobulid genomes as the focus, with the two outgroups linked by a direct pairwise alignment (*L. erinacea* – *M. hypostoma* – *M. yarae* – *H. sabinus*). Chromosomes are referred to by a two-letter species prefix (MH, MY, HS, LE) followed by their assembly designation corresponding to the numbers shown in Figure 2B.

**Figure 2.**
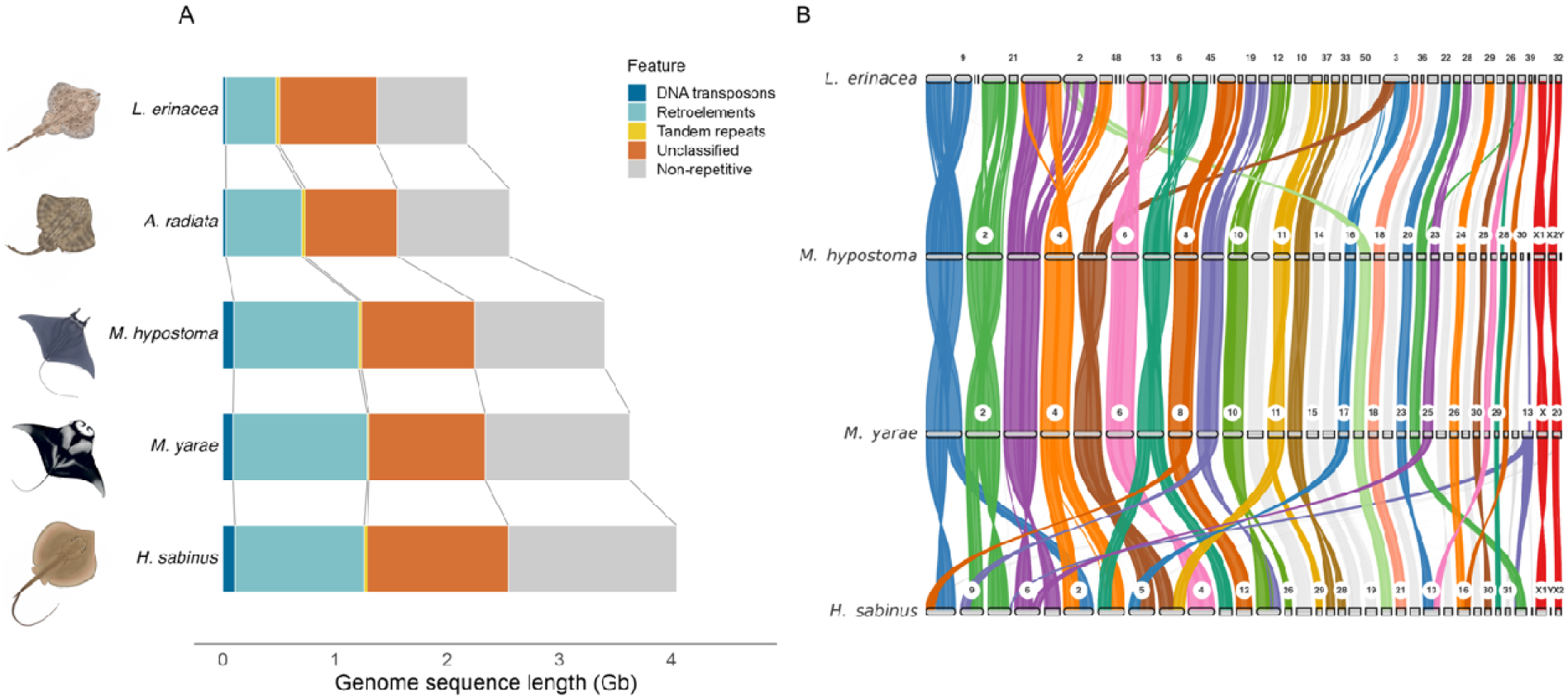
Repeat content and chromosome-scale synteny across batoid genomes. (A) Repeat composition of the five batoid assemblies, ordered by assembly size. Bars show the total length (Gb) of DNA transposons, retroelements (LINEs, LTR elements, SINEs), tandem repeats (satellites and simple repeats), unclassified repeats and non-repetitive sequence, as identified by RepeatMasker. Grey lines link equivalent category boundaries between species. Species illustrations copyright Marc Dando. (B) Genome-wide synteny derived from the Cactus alignment. Ribbons connect syntenic blocks between *L. erinacea* and *M. hypostoma*, *M. hypostoma* and *M. yarae*, and *M. yarae* and *H. sabinus*. *Mobula hypostoma* chromosomes are ordered by size, with sex chromosomes on the right. Chromosomes of all other species are ordered by the position of their principal homolog in the adjacent track. Ribbons are coloured when the alignment between a chromosome pair accounts for at least 15% of the aligned sequence of each chromosome and the pair is involved in a fusion or fission event in at least one comparison. Colours are assigned by the corresponding *M. hypostoma* chromosome. All other alignments including one-to-one homologies and alignments below the 15% threshold are shown in grey. Sex chromosomes are shown in red. Chromosome numbers follow those designated in each assembly and for clarity, only every second chromosome is labelled along with all sex chromosomes. Chromosome identifiers used in the text (e.g. MH14) combine a species prefix with the chromosome numbers shown.

### Structural genome rearrangements during batoid evolution

We quantified inferred structural rearrangements (SR) along our five species batoid phylogeny leveraging the ancestral genomes reconstructed by Cactus. SRs were extracted from the HAL alignment using *halSummarizeMutations* (*--maxNFraction* 0), which classifies rearrangements into insertions, inversions, transpositions, duplications and deletions. For each branch, SRs were identified by comparing each descendant genome to its immediate ancestor, enabling the assignment of rearrangements to particular evolutionary lineages. We focussed on myliobatiform evolution by summarising SRs that occurred along the branches from Anc2 to Anc3, Anc2 to *H. sabinus* and Anc3 to *M. hypostoma* and *M. yarae* (Figure 3A).

**Figure 3.**
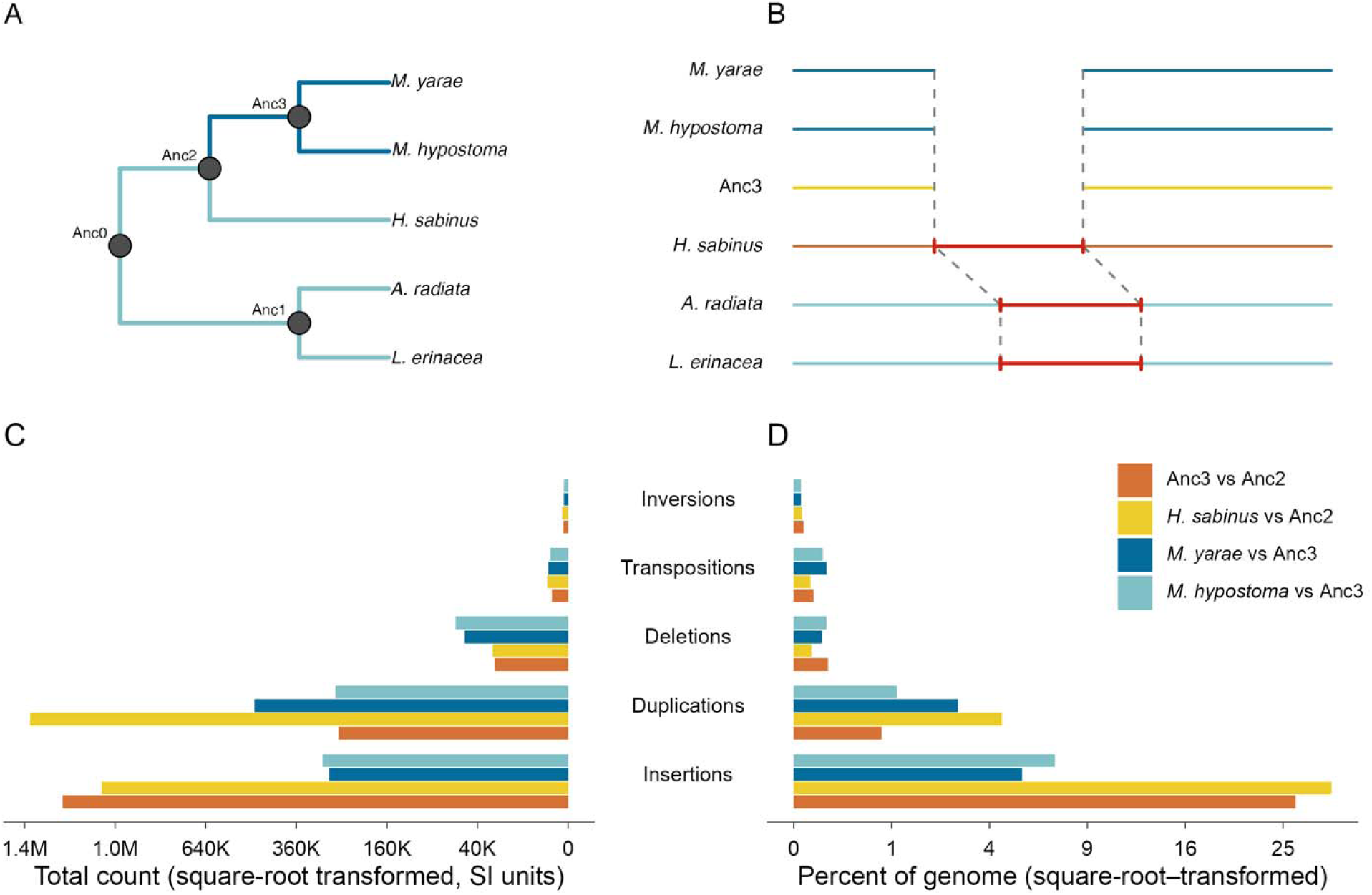
(A) Cladogram depicting phylogenetic relationships among the five target batoid species and their reconstructed ancestors. Pelagic species are shown in dark blue and benthic species in light blue. Terminal branches represent extant species, while internal nodes correspond to ancestral genomes reconstructed using Cactus: Anc0 (ancestral genome for all species); Anc1 (common ancestor of little skate and thorny skate); Anc2 (common ancestor of Atlantic stingray and mobulids); and Anc3 (common ancestor of mobulids). (B) Schematic representation of the mobulid specific deletions targeted in this study. Each horizontal line represents a syntenic genomic region with red bars indicating sequence presence and gaps representing lineage-specific deletions. We identified deletions inferred to have occurred between Anc2 and Anc3 (and therefore absent in Anc3, *M. hypostoma* and *M. yarae).* (C–D) Bar plots summarising five classes of SRs (insertions, duplications, deletions, transpositions and inversions) identified through whole genome alignment. Rearrangements were detected between *M. hypostoma*, *M. yarae*, *H. sabinus* and their inferred common ancestor, as well as between reconstructed ancestral genomes Anc2 and Anc3. (C) Total counts of structural rearrangement class. (D) Proportion of each genome affected by each SR class.

### Functional annotation of ancestral mobulid deletions

To identify candidate regions associated with mobulid adaptation, we functionally annotated the deletions inferred to have occurred along the branch between Anc2 and Anc3 (Figure 3A-B). We focused on deletions because their impact on annotated gene features could be readily inferred, which was challenging for other SR types. Since no genome annotation exists for Anc2, we used the annotated genome of *H. sabinus* as a proxy to infer gene content in the ancestral genome. To do this, the deletion coordinates from the *halSummarizeMutations* output (Anc2 vs Anc3) were formatted into BED files and expanded by 1 bp on each side to create a flanking region so that the resulting subalignments captured the deletion breakpoints in all genomes. We then converted our HAL file into a MAF using hal2maf (Hickey et al., 2013), from which we extracted the deletion intervals using *mafExtractor* from MafTools (Earl et al., 2014). We then used a custom R script to parse the resulting MAF subalignment into a summary table, capturing locations for each deletion and flanking sequences in the *H. sabinus* genome, enabling downstream functional annotation. Deletions were retained for annotation only where the deleted sequence was present in all three outgroup genomes (*H. sabinus*, *L. erinacea* and *A. radiata)*, thereby targeting ancestral batoid sequence that was lost in the mobulid ancestor. To functionally annotate these regions, genomic coordinates were formatted into BED files and analysed with SnpEff v5.2c (Cingolani et al., 2012). As a SnpEff database does not exist for *H. sabinus*, we built a custom database using the *H. sabinus* annotation (GCF_030144855.1_sHypSab1.hap1_genomic.gtf.gz) and SnpEff’s build command. SnpEff was then run to identify the functional context of each deletion (intronic, exonic or intergenic) and annotate regions overlapping known genes in the *H. sabinus* database. To identify candidate genes underlying mobulid-specific morphological traits, we screened annotations for those affecting genes involved in craniofacial, oral and limb development. Genes were prioritised based on their association with three Gene Ontology (GO) terms: “cranial skeletal system development” (GO:1904888), “roof of mouth development” (GO:0060021) and “limb development” (GO:0060173), as described in the Mouse Genome Informatics (MGI) database (Table S3). Additional genes were identified from the literature on vertebrate craniofacial development (Lamichhaney et al., 2015; Mork & Crump, 2015, Table S4).

### Gene Ontology enrichment analyses

Gene Ontology (GO) enrichment analysis was conducted to identify biological processes overrepresented among genes affected by regions present in Anc2 but absent in the mobulid ancestor (Anc3). Gene names and symbols were extracted from the BED files containing deletion coordinates. Gene IDs were downloaded from NCBI and matched to gene names using a custom R script to obtain the input files for GO analysis. The *H. sabinus* GO annotation was downloaded from NCBI (GCF_030144855.1-RS_2023_09_gene_ontology.gaf.gz) and used to create a custom *H. sabinus* annotation with the AnnotationForge v.1.46.0 R package (Carlson & Pagès, 2025). GO enrichment of Biological Process terms was performed in R v4.4.1 using the *enrichGO* function from clusterProfiler v4.12.6 (Wu et al., 2021) with the keytype set as “GID” and an adjusted P-value (Benjamini-Hochberg) threshold of 0.1. The gene set from the BED file (2,413 genes, of which 1,950 had GO annotations) was tested against the full set of GO-annotated genes (14,303) as the background. For visualisation, redundant significantly enriched GO terms were collapsed using the *simplify* function from clusterProfiler with a similarity cutoff of 0.7, retaining the most significant term for each cluster.

## Results

### Genome assembly and quality assessment

We generated chromosome-level reference genomes for *M. hypostoma* and *M. yarae* using a combination of long- and short-read sequencing together with Omni-C chromatin conformation capture data. The *M. hypostoma* assembly is a haploid representation of the genome (sMobHyp1.1) and has a total length of 3.48 Gb, comprising 31 autosomes, one Y and two X chromosomes (X1 and X2). Contig and scaffold N50 values were 23.3 Mb and 152.4 Mb, respectively (Table 1, Table S1 and Figure 1C–D). Quality assessment showed no contamination, a consensus accuracy of QV49 and assembly BUSCO completeness of 93.4% [S:91.0%, D:2.4%], F:2.5%, M:4.1% (n:3354, vertebrata_odb10). NCBI RefSeq annotation identified 44,060 mRNAs from 18,916 protein-coding and 3,892 non-coding genes (GCF_963921235.1-RS_2024_02). BUSCO completeness for the annotation was 97.3% [S:94.7%, D:2.7%], F:0.6%, M:2.1% (n:3354, vertebrata_odb10). The mitogenome was successfully assembled into a circular sequence of 18,142 bp in length. Notably, *M. hypostoma* represents the first cartilaginous fish assembly leveraging ONT sequencing as the base technology, with most current studies using PacBio HiFi reads (Figure S4).

The *M. yarae* assembly pipeline generated two complete, haplotype resolved assemblies (sMobBir1.hap1 and sMobBir1.hap2), each comprising 32 autosomes and one X chromosome (X). However, our synteny analysis indicates that chromosome 20 corresponds to *M. hypostoma* X2, suggesting that *M. yarae* also possesses two X chromosomes (see Synteny comparisons across batoid genomes for further details). The sMobBir1.hap1 assembly had a total length of 4.01 Gb, contig and scaffold N50 values of 7.3 Mb and 148.7 Mb respectively, a total gap length of 30.85 Mb, and a consensus accuracy of QV61 (Figure S5 A–B). Assembly BUSCO completeness was 93.0% [S:90.8%, D:2.2%], F:3.4%, M:3.7% (n:3354, vertebrata_odb10). The sMobBir1.hap2 assembly had a total length of 3.63 Gb, contig and scaffold N50 values of 8.6 Mb and 152.4 Mb, respectively, a total gap length of 54.98 Mb, and a consensus accuracy of QV63 (Figure S5 C–D). Assembly BUSCO completeness was 91.2% [S:89.1%, D:2.1%], F:3.5%, M:5.3% (n:3354, vertebrata_odb10). Gene annotation was carried out on sMobBir1.hap1, yielding 45,885 mRNAs from 20,172 protein-coding and 6,132 non-coding genes (GCF_030028105.1-RS_2025_03). BUSCO completeness for the annotation was 96.8% [S:94.0%, D:2.8%], F:1.0%, M:2.2% (n:3354, vertebrata_odb10).

Telomeric repeat signal was identified in the *M. hypostoma* assembly and in both haplotype assemblies of *M. yarae*. Telomeric repeats were detected at one or both ends of 8/34 (24%) scaffolds >= 5Mb in *M. hypostoma*, 13/33 (39%) in sMobBir1.hap1 and 9/33 (27%) in sMobBir1.hap2 (Figure S6). Several chromosomes had intact telomeric repeats at both ends including four large chromosomes in *M. hypostoma* and the X1 chromosome of sMobBir1.hap1 (Figure S6). These results are consistent with the high contiguity metrics reported for the underlying assemblies.

### Repeat landscape

Repetitive sequences accounted for between 61–66% of each batoid genome, with *M. hypostoma* showing the highest repeat content (Figure 2A, Table S2). Retroelements dominated across all species, particularly LINEs, which were most abundant in the mobulids (*M. hypostoma*: 29.28%; *M. yarae*: 29.64%). The predominant LINE family in all batoids was L2/CR1/Rex, which accounted for between 12.5–22% of genome content. Long terminal repeats (LTRs) contributed to 2.7–4.7% of genome content, with Gypsy/DIRS1 elements predominating (2.5–3.7%) and retroviral LTRs making up only a small fraction (0.15–0.63%). DNA transposons made up a small fraction of each batoid genome, with *M. hypostoma* having the highest (2.90%) and *A. radiata* the lowest (0.93%) representation of these elements (Figure 2A).

### Synteny comparisons across batoid genomes

To uncover conserved and rearranged features of batoid genome architecture, we examined synteny patterns leveraging our multi-species Cactus alignment. Strongest synteny was observed between *M. hypostoma* and *M. yarae*, with near complete one-to-one homology across chromosomes (Figure 2B). Specifically, for 33 of the 34 *M. hypostoma* chromosomes, over 99.8% of the aligned sequence matched a single *M. yarae* chromosome, with 24 chromosomes aligning exclusively to a single *M. yarae* chromosome. One notable exception was for *M. yarae* chromosome MY13, for which only ∼4% aligned to *M. hypostoma*, mostly to MH31. Interestingly, *M. hypostoma* X2 showed one-to-one homology with *M. yarae* chromosome 20 (MY20), with 99.9% of aligned MHX2 sequence mapping to MY20. This suggests that although designated as an autosome, MY20 likely represents the X2 chromosome in *M. yarae*. The *M. hypostoma* Y chromosome (MHY) displayed minimal alignment to both MY20 and MYX – as expected given the absence of a Y chromosome in the *M. yarae* assembly – while MHX1 showed near-complete one-to-one homology with MYX.

Comparison of *M. yarae* and *H. sabinus* revealed a mix of both conserved and rearranged chromosomes (Figure 2B). Thirteen of the 31 *M. yarae* autosomes showed reciprocal one-to-one homology with *H. sabinus*, particularly among the smaller chromosomes (e.g. MY15 and HS17; MY19 and HS20; MY27 and HS27; MY29 and HS24, MY30 and HS30). A further six *M. yarae* chromosomes each corresponded to part of a single *H. sabinus* chromosome – consistent with fission – for example, HS13 aligned to non-overlapping regions of MY23 and MY31; and HS16 with MY26 and MY32. Conversely, ten large *M. yarae* chromosomes comprised large blocks homologous to two *H. sabinus* chromosomes – consistent with fusion – for example MY3 comprises HS14 and part of HS6, MY7 comprises HS15 and part of HS3, MY8 comprises HS12 and part of HS1, and MY14 comprises HS25 and HS28. Sex chromosome synteny was largely retained between the two species with MYX and MY20 (X2) corresponding predominantly to HSX1 and HSX2, respectively.

Synteny comparison between *M. hypostoma* and *L. erinacea* revealed a similar number of one-to-one homologies to the *H. sabinus* comparison but the larger number of *L. erinacea* chromosomes was reflected by extensive fusion (Figure 2B). Fourteen *M. hypostoma* chromosomes displayed synteny with two or more *L. erinacea* chromosomes, seven of which correspond to two complete *L. erinacea* chromosomes (e.g. MH1 comprises LE4 and LE9; MH11 comprises LE18 and LE37). In contrast, several *L. erinacea* chromosomes aligned to two *M. hypostoma* chromosomes – consistent with fission – for example LE3 aligns to both MH5 and MH16, and LE26 to both MH26 and MH28. *M. hypostoma* sex chromosome MHX1 corresponds to two complete *L. erinacea* chromosomes (LE27 and LE40), whereas MHX2 shows one-to-one homology with LE32. Taken together, these findings suggest that karyotype evolution in mobulids has involved both the maintenance of many ancestral chromosomes as well as several large-scale fission and fusion events.

### Structural genome rearrangements during batoid evolution

To investigate patterns of genome evolution in Batoidea, we traced the distribution of structural rearrangements (SRs) across the batoid phylogeny by leveraging ancestral genomes reconstructed by Cactus. SRs were characterised by comparing each descendant node to its immediate ancestor, allowing us to map rearrangements to specific branches of the tree (Figure 3A–B). We focussed on SRs that occurred during myliobatiform genome evolution. Structural rearrangements were widespread across the myliobatiform tree yet displayed notable differences in frequency and composition (Figure 3C–D). Along the ancestral mobulid branch (Anc3), approximately 1.5 million SRs were identified, affecting 27% of the Anc3 genome. In contrast, the *H. sabinus* lineage accumulated 2.5 million SRs, impacting 35% of its genome. These proportions reflect the fraction of each genome that has been structurally rearranged along each respective branch (Figure 3D).

Insertions were the most common SR category across all branches, comprising up to 81% of all rearrangement events and accounting for the majority of rearranged sequences (64–96%, Figure 3C–D). The mobulid ancestor (Anc3) gained 849 Mb of inserted sequence (26% of its genome), compared to 1.2 Gb in *H. sabinus* (30% of its genome). Among extant mobulids, *M. hypostoma* accumulated more insertions than *M. yarae* (247 Mb vs 197 Mb), representing 7% and 5% of their respective genomes (Figure 3C–D). Duplications were also frequent, representing 17–59% of SRs across the myliobatiform branches. The mobulid ancestor (Anc3) accumulated ∼257,000 duplications (Figure 3C), while 1.1% and 2.8% of *M. hypostoma* and *M. yarae* genomes can be attributed to duplications, respectively (Figure 3D).

Deletions were among the most infrequent class of SR across myliobatiform lineages (Figure 3C–D). Along the ancestral mobulid branch (Anc3 vs Anc2), approximately 26,000 deletions were identified, representing 1.7% of all inferred SRs. Interestingly, this was similar to the number identified along the much longer *H. sabinus* branch (∼27,600), but encompassed three times as much deleted sequence (3.9 vs 1.3 Mb). The extant mobulids showed similar numbers of deletions since their separation (*M. hypostoma* vs. Anc3 and *M. yarae* vs Anc3), with 61,000 and 52,000 inferred in *M. hypostoma* and *M. yarae*, respectively. Overall, deletions represented less than 1% of total genome sequence across all branches examined (Figure 3C–D). Transpositions and inversions were also relatively minor contributors to overall genome rearrangements, accounting for <1% of all SRs and affecting ∼0.1% of genome sequence per branch (Figure 3C–D).

### Functional impact of ancestral mobulid deletions

To investigate genomic signatures associated with the transition to a pelagic lifestyle, we annotated deletions occurring along the ancestral mobulid branch (Anc3 vs Anc2, Figure 3B). Deletions were targeted owing to: i) our observation that the inferred rate of deletion occurrence appeared markedly elevated at the base of mobulid evolution (see above) compared to the sister stingray lineage, ii) past observations that deletions can modify developmental phenotypes by altering the context-specific expression of master regulatory genes via alteration/ablation of regulatory elements (Chan et al., 2010; Thompson et al., 2018), including in association with major body-plan transitions (Kvon et al., 2016) and iii) because it was possible to more readily infer their functional impacts compared to other SR types, leveraging the stingray reference genome, which by definition retains the deleted regions.

We tested the hypothesis that ancestral mobulid deletions disrupted non-coding regions in vertebrate genes with known roles controlling development processes such as cranial skeletal system development and roof of mouth development (Table S3-4), facilitating the evolution of devil ray morphological synapomorphies. Of the ∼26,000 deletions inferred along the ancestral mobulid Anc3 branch, 7,059 involved sequence present in all three outgroup genomes and were retained for annotation. Of these, 122 were associated with predicted orthologs for 68 pre-defined genes. Of these, 53 occurred within intronic regions of 39 affected genes (Tables S3-4). In contrast, nine deletions occurred in exonic sequences or regions immediately upstream of the first exon of seven affected genes, potentially affecting proximal cis-regulatory elements (Figure 4A-D). For example, we identified a deletion approximately 1 kb upstream of the first exon of *shh* – encoding sonic hedgehog – a crucial signaling molecule for vertebrate craniofacial development (Dworkin et al., 2016; Figure 4A). Similar upstream deletions were detected near the first exons of *alx3* and *fgfr3* (Figure 4B; Table S4), genes essential during craniofacial differentiation and skeletal development, respectively (Lakhwani et al., 2010; Mitchell et al., 2021; Wilke et al., 1997). Deletions also affected non-coding exons in *bpnt2* and *mthfd1l*, both linked to craniofacial and neural tube development defects (Dubail & Cormier-Daire, 2021; Momb et al., 2013; Figure 4C-D).

**Figure 4.**
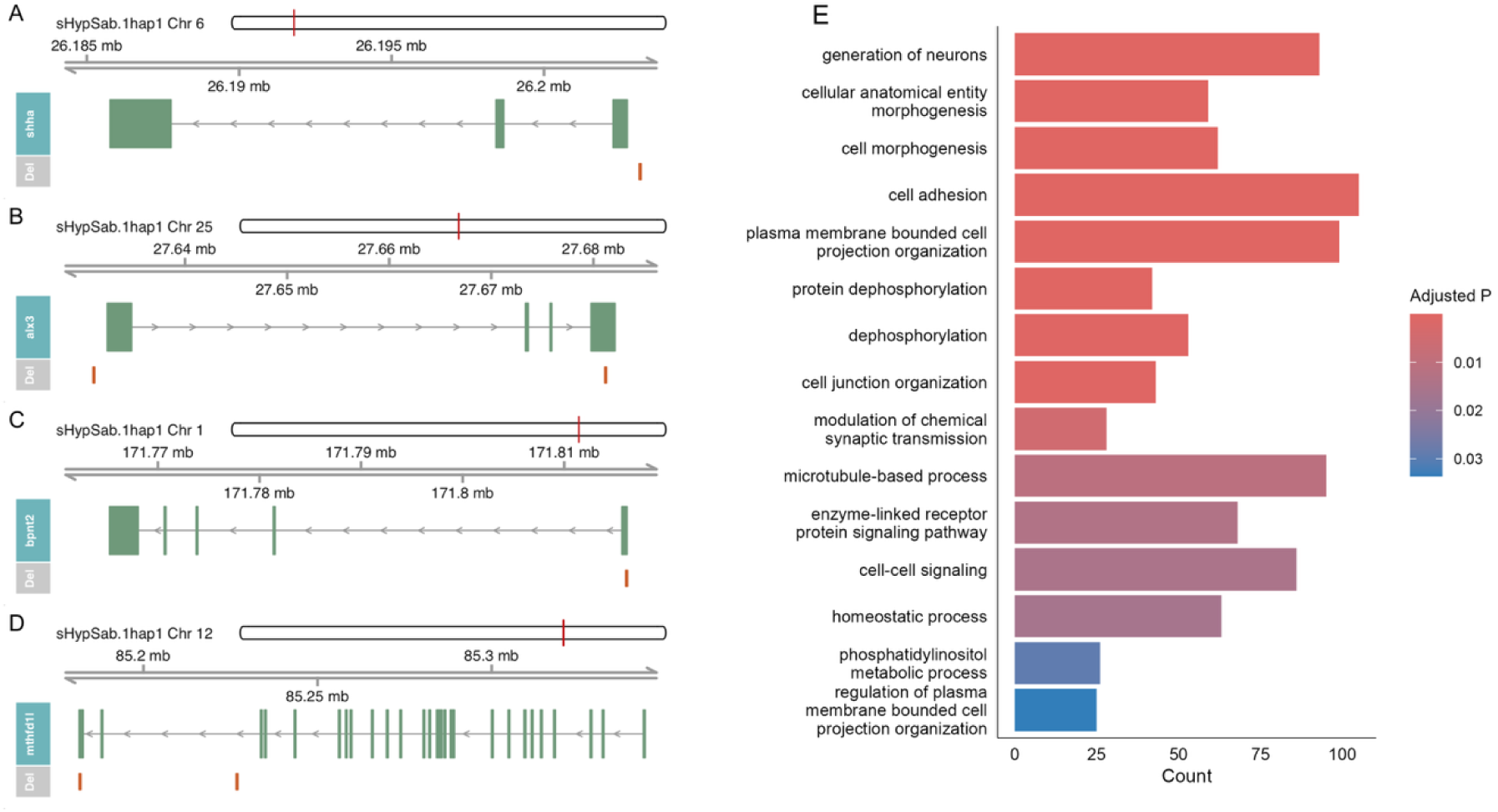
Gene track visualisation for four of the seven genes in which ancestral mobulid deletions were identified within exonic sequences or regions immediately upstream of the first exon: *shha* (A), *alx3 (B)*, *bpnt2* (C) and *mthfd1l* (D). Deleted regions were originally identified in the Anc2 genome but annotated with respect to their orthologous positions in the *H. sabinus* assembly (sHypSab1). The chromosomal location of each gene is indicated by a red line in the top ideogram track. Genomic coordinates are displayed along the axis track. Exons are represented by green boxes with directional arrows showing the strand orientation. Deletions are shown in orange along the bottom track. (E) Gene Ontology (GO) Biological Process terms significantly enriched among genes affected by ancestral mobulid-specific deletions (adjusted P ≤ 0.1). Redundant terms were collapsed based on semantic similarity. The 15 most significant terms are shown and ordered by significance. The x-axis shows the number of genes associated with each term and the colour of the bars represent adjusted p-values with pink shades indicating stronger enrichment. See Table S5 for the full list.

To formally test if deletions were significantly enriched for terms relating to morphological or craniofacial development, we conducted GO analysis using 14,303 annotated *H. sabinus* genes as a genomic background. We identified 61 significantly enriched GO terms, with neurogenesis and nervous system development being the most prevalent, accounting for 24 terms overall (Figure 4E and Table S5). The most enriched terms included “generation of neurons” (GO:0048699; adjusted p-value <0.0001); “neurogenesis” (GO:0022008; adjusted p-value <0.0001); “neuron development” (GO:0048666; adjusted p-value <0.0001); “neuron differentiation” (GO:0030182; adjusted p-value <0.0001); and “neuron projection development” (GO:0031175; adjusted p-value <0.0001). A substantial overlap in genes contributed to these GO terms, including *lhx6a*, *sema3h*, *plxnd1* and *crtac1b*. Additionally, some genes associated with neurogenesis overlapped with the broader development term “regulation of developmental process” (GO:0050793, adjusted p-value = 0.087), underscoring their potential developmental significance. Overlapping genes between these terms included *sema3h*, *plxnd1*, *pacsin1b*, *sema6dl*, and *runx2a*. These functional enrichments are consistent with the individual-gene level annotations identified using SnpEff.

## Discussion

A core achievement of this study is the generation and sharing of chromosomal-level reference genomes for *M. hypostoma* and *M. yarae*, both annotated using extensive transcriptomic data. Elasmobranch genomes are notoriously difficult to assemble due to their large size, high repeat content and complex structural features (Nishimura et al., 2022). The difference in contiguity between the reported assemblies, despite comparable functional completeness scores, is likely due to the use of Oxford Nanopore Technology (ONT) in *M. hypostoma.* Although the majority of *M. hypostoma* ONT reads were <50 kb, we nonetheless captured a large number of ultra-long reads (including >100 kb), which likely aided in spanning complex repeat structures during contig assembly. In contrast, the *M. yarae* assembly, which used PacBio HiFi reads, achieved lower contiguity (contig N50: 7.30 Mb) despite comparable BUSCO completeness, highlighting the advantages of ONT technologies for resolving complex elasmobranch genome architectures. These resources not only set benchmarks for genome quality in chondrichthyans but provide platforms for investigating mobulid evolution and conducting high-resolution comparative studies across sharks and rays.

### Karyotype reorganisation and sex chromosome evolution in batoids

Our whole genome alignments revealed a combination of both conserved and rearranged chromosomes between the mobulids and both *H. sabinus* and *L. erinacea,* while genomic architecture remained almost entirely conserved between the two mobulid genomes. Our observations are consistent with a scenario of substantial karyotypic reorganisation in the mobulid ancestor, followed by minimal chromosomal evolution within mobulids, and also reflect the closer relationship of the two mobulids, which are thought to have diverged approximately 20 Mya (Poortvliet et al., 2015). While many smaller *M. yarae* chromosomes retain one-to-one homology with *H. sabinus*, most of the largest *M. yarae* chromosomes appear to originate from fusions of distinct, non-overlapping *H. sabinus* chromosome regions. This suggests that the current *H. sabinus* karyotype may more closely reflect an ancestral myliobatiform state, with a limited number of large-scale chromosomal fusions characterising mobulid genome evolution. This is consistent with early cytogenetic work attributing the large meta- and submetacentric chromosomes of *M. mobular* (reported as *M. japonica*) to centric fusions (Asahida et al., 1993). Further reconstruction of ancestral myliobatiform and mobulid karyotypes together with combined synteny analyses between rajiform and myliobatiform genomes will be needed to further explore this and alternative scenarios. Chromosome rearrangements are widely recognised as catalysts of speciation and adaptation (Feulner & De-Kayne, 2017; Guerrero & Kirkpatrick, 2014; Wellenreuther et al., 2019). In particular, chromosome fusions can bring together previously unlinked loci, creating novel, co-evolving gene clusters with reduced recombination rates (de Vos et al., 2020; Liu et al., 2022; Yoshida et al., 2023). It is possible that mobulid-specific chromosomal fusions may have combined gene groups advantageous for a highly mobile, pelagic lifestyle. Further studies are necessary to ascertain whether these putatively fused chromosomes are enriched for loci underpinning mobulid-specific adaptive traits.

Sex chromosome composition in *M. hypostoma* mirrored that of *H. sabinus*, comprising two X chromosomes and one Y (X1X2Y). In *M. yarae*, only a single X chromosome was formally identified during manual curation; however, sequences corresponding to *M. hypostoma* X2 mapped almost exclusively to the chromosome-length scaffold designated as chromosome 20. This suggests that *M. yarae* also has an X1X2Y system, with chromosome 20 representing the X2. This was likely overlooked during curation because a multiple sex chromosome system was not anticipated. The presence of this configuration in both Mobula species and in *H. sabinus* suggests that X1X2Y represents the ancestral myliobatiform state. This interpretation is consistent with recent comparative genomic studies across elasmobranchs, which report the emergence of an X1X2Y system in batoids through fusion of the ancestral Y with an autosome, followed by differentiation of a neo-X chromosome (Lee et al., 2025; Niwa et al., 2025). In line with this, *M. hypostoma* X2 showed one-to-one homology with a single *L. erinacea* chromosome, consistent with X2 deriving from an ancestral autosome. Under an X1X2Y system and assuming 31 autosomes, both mobulid genome assemblies predict a diploid number of 2n = 66 in females and 2n = 65 in males. The only published mobulid karyotype reported 2n = 66 for both an adult female and a male foetus (Asahida et al. 1993). This is broadly consistent with our inferred chromosome complements, however the reported count in males differs by one chromosome. Notably, Asahida et al., (1993) observed morphological differences in one chromosome pair between the sexes, suggesting possible sex chromosome differentiation, yet were unable to resolve the underlying chromosome configuration. Further comparative work across a broader range of batoid genomes, together with cytogenetic analysis of both sexes, will clarify when and how the X1X2Y system emerged and diversified in rays.

### Repeat content and genome size variation in batoids

Elasmobranch genomes vary considerably in size, largely attributable to differences in repeat content (Kuraku et al., 2025). To explore this relationship in batoids, we characterised repeat composition across five batoid species, with genome sizes ranging from 2.2 to 4.0 Gb. We found that repetitive sequences make up over 60% of each batoid genome, exceeding typical values reported in teleosts and tetrapods, but in line with recently reported values in the red stingray (61.2%) and whitebelly skate (63.9%) (Kuraku et al., 2025). Interestingly, both mobulid species displayed comparable levels of non-repetitive genomic content, suggesting that genome size variation observed between *M. hypostoma* and *M. yarae* is driven primarily by repetitive DNA. Furthermore, the comparable abundance of retroelements in both mobulid genomes and *H. sabinus* suggests a lineage-wide repeat expansion in the ancestral myliobatiform clade. Within mobulids, retroelements – particularly LINEs – dominated the repetitive sequence landscape, a pattern consistent with observations in other elasmobranch species including the brownbanded bamboo shark, whale shark and cloudy catshark (Hara et al., 2018). Retroelements are known drivers of genomic innovation and structural rearrangements, capable of disrupting coding and regulatory sequences, modifying promoters and generating novel exons (Böhne et al., 2008; Bourque, 2009). In line with this, proliferation of repetitive sequences has been associated with the origin of genes, for example during the emergence of placental protein-coding genes in mammals (Blaise et al., 2003). Our findings are consistent with the idea that repeat mediated structural rearrangements potentially played a role in shaping the ancestral mobulid genome, but additional work will be required to explore this hypothesis.

### Structural rearrangements as potential drivers of mobulid adaptation

To investigate candidate genomic changes associated with ancestral mobulid adaptations, we characterised the distribution of SRs across myliobatiform genomes, focussing on changes occurring in the common mobulid ancestor after divergence from the stingray lineage. Insertions and duplications were the dominant SR type, accounting for over a quarter of the reconstructed ancestral mobulid genome (Anc3) length. While most of these SRs are likely to be selectively neutral, a subset may have altered gene function or regulation (McGee et al., 2020). Inversions and transpositions, although rare, affected over one million base pairs of the ancestral mobulid genome, highlighting their potential significance, as such rearrangements can suppress recombination between diverging lineages and facilitate reproductive isolation and adaptation (Knief et al., 2016; Pei et al., 2025). Further functional characterisation of genes within these rearranged regions may help clarify their contribution to mobulid-specific adaptations.

### Candidate deletions associated with mobulid morphological evolution

The vast majority of ancestral mobulid-specific deletions found to associate with our prioritised gene set were located in intronic or intergenic regions. Non-coding *cis*-regulatory sequences such as enhancers play a central role in morphological evolution and divergence (Onimaru, 2020; Rajderkar et al., 2024; Rebeiz & Tsiantis, 2017). For example, disruption to enhancer elements proximal to craniofacial genes has been shown to cause skeletal defects in mouse models (Crane-Smith et al., 2023), and differences in enhancer landscape has been implicated in the craniofacial divergence between humans and chimpanzees, despite a background of highly conserved coding sequence (Prescott et al., 2015). It is therefore plausible that intergenic or intronic deletions near genes regulating craniofacial development disrupted enhancer sequences, thereby contributing to differential gene expression and the emergence of derived head morphology in mobulids.

Several deletions overlapped the putative promoter or untranslated regions (UTRs) of key developmental signalling and regulatory genes, including sonic hedgehog (Shh) and fibroblast growth factor (FGF) signalling pathways. Given the role of promoter architecture in regulating transcription (Tian et al., 2025), such deletions may have altered gene expression levels during embryonic craniofacial development. In particular, *shha*, which is expressed in the frontonasal domain and controls cranial skeletal formation and tissue differentiation (Adameyko & Fried, 2016), is a compelling candidate for mobulid facial remodelling. A deletion in the 5’ UTR of *fgfr3* – a gene involved in cranial cartilage formation (Adameyko & Fried, 2016; Morice et al., 2023) – provides additional support for functional disruption of craniofacial architecture. Furthermore, a deletion overlapping the putative promoter of *alx3,* an ALX homeobox gene, responsible for regulating frontonasal cell differentiation during embryonic development (Lakhwani et al., 2010; Mitchell et al., 2021), may also contribute to the disturbance of key developmental pathways. Taken together, these preliminary findings lead us to speculate that a polygenic architecture underlies batoid craniofacial developmental evolution, involving collective disruption of regulatory elements and expression domains. Future functional and developmental datasets and analyses will be required to fully explore this hypothesis.

Genes affected by mobulid-specific deletions were also significantly enriched for biological pathways associated with neurogenesis and nervous system development. Several of these genes play crucial roles in vertebrate embryogenesis. For example, the transcription factor *runx2* – a major regulator of skeletal development in Osteichthyes – has been shown to influence dorso-ventral patterning in zebrafish embryos (Flores et al., 2008). While the function of *runx2* in cartilaginous fish may differ from its bony vertebrate orthologs, its involvement in the formation of cartilaginous structures including the elasmobranch chondrocranium is plausible. Deletions affecting regulators like *runx2* may therefore have contributed to craniofacial divergence in the mobulid lineage. Emerging evidence suggests that neurogenesis and craniofacial developmental pathways are closely linked (Adameyko & Fried, 2016). For example, early craniofacial patterning and differentiation is orchestrated by migrating neural crest cells that populate the fronto-nasal region and initiate the formation of facial structure (Adameyko & Fried, 2016; Dupin et al., 2006). Beyond this initial stage, signals from the developing brain and peripheral nervous system continue to shape craniofacial architecture through pathways such as *ssh* (Adameyko & Fried, 2016). The enrichment of neurogenesis-related deletions in mobulids therefore supports the notion that nervous system and craniofacial development are mechanistically linked. Further work to directly infer the functional effects of mobulid specific deletions will shed light on the drivers of mobulid adaptation.

## Conclusion

The evolutionary scenarios described above are consistent with a transition from a benthic, durophagous ancestor – likely resembling modern cownose rays (*Rhinoptera) –* to a pelagic and planktivorous lifestyle in mobulids. Rhinopterids possess several features also present in mobulids including wing-like pectoral fins and small, fused cephalic lobes which are ventrally located and used for feeding (Swenson et al., 2018). These shared traits suggest that the most recent common ancestor of rhinopterids and mobulids already possessed structural adaptations that facilitated ecological niche expansion into open water. A shift from ventral to forward or sub-terminal mouth orientation likely enabled the transition from benthic to pelagic feeding, subsequently leading to niche differentiation and reproductive isolation from other myliobatiforms. This ecological shift may have been reinforced by lineage-specific adaptations, including the development of cephalic lobes (Swenson et al., 2018), that led to the distinctive mobulid phenotype we know today. Gaining functional support for this scenario will ultimately require a reference genome for a cownose ray, enabling further comparative genomic analyses of mobulid and related myliobatiform or batoid members, alongside comparative transcriptomic and regulatory element datasets that inform the basis for variation in developmental phenotypes shaping craniofacial morphology and other mobulid adaptations.

## Acknowledgements

Sequencing and assembly of the *M. hypostoma* genome was funded by a Royal (Dick) School of Veterinary Studies and Roslin Institute pump priming grant awarded to EH and DJM. We would like to acknowledge Trey Knott at NOAA and Tomàs Marquès, Alba Duch Larregola and Esther Lizano at Universitat Pompeu Fabra who were instrumental in facilitating the shipment of *M. hypostoma* blood samples from Florida for sequencing. We are grateful to the Save Our Seas Foundation and the Mote Scientific Foundation for funding the devil ray fieldwork in Florida. We also thank the Mote Marine Laboratory’s Marine Operations team (P. Hull, D. Dougherty, G. Byrd and C. Cole), the Destin-Fort Walton Beach Natural Resources team (A. Fogg, J. Valek, M. Norberg) and several field operations graduate students and interns who assisted with collecting valuable samples for this study.

## Data availability statement

Sequencing data and genome assemblies generated in this study have been made available on public databases. For *Mobula hypostoma*, data are collected under the umbrella BioProject PRJEB71674 at the European Nucleotide Archive, comprising raw Oxford Nanopore long-read, Hi-C, Illumina short-read sequencing data (PRJEB71672) and the chromosome-level genome assembly sMobHyp1.1 (PRJEB71673; GenBank accession GCA_963921235.1). RNA-seq reads are available from the European Nucleotide Archive under study accession PRJEB71148. For *Mobula yarae*, all data are collected under the umbrella BioProject PRJNA954412 at NCBI, comprising raw PacBio HiFi and Hi-C sequencing data (PRJNA1218824), RNA-seq data (PRJNA1218823) and the chromosome-level genome assemblies sMobBir1.hap1 (PRJNA924117; GenBank accession GCA_030028105.1) and sMobBir1.hap2 (PRJNA924118; GenBank accession GCA_030035685.1). *Mobula* yarae genome assemblies were generated and deposited under the original species designation and therefore retain *M. cf. birostris*-based identifiers (e.g. sMobBir1). Code used for analysis is available on GitHub (https://github.com/lsoler-clavel/mobulid-comp-genomics-craniofacial-dev) and archived on Zenodo (Soler Clavel, 2026).

## Benefit sharing statement

Benefits from this research accrue from the sharing of our data on public databases as described above.

## Author contributions

EH and DJM designed research and provided supervision to LSC; ABP, KBH and LH collected samples; GN, EJ and SHL contributed materials and resources; MG sequenced the *Mobula hypostoma* genome; JGG, FC and TSA assembled the *Mobula hypostoma* genome; TT, NJ and JB sequenced the *Mobula yarae* genome; CW sequenced the *Mobula yarae* transcriptome; DEA, LA, OF, BF and GF assembled the *Mobula yarae* genome. LSC performed the cactus alignment and downstream analysis with input from MKG; LSC and EH wrote the paper with input from all coauthors.

## Notes

### Competing Interest Statement

The authors have declared no competing interest.

