## Supplementary Figures for "Manta and devil ray genome assemblies uncover signatures of pelagic adaptation"

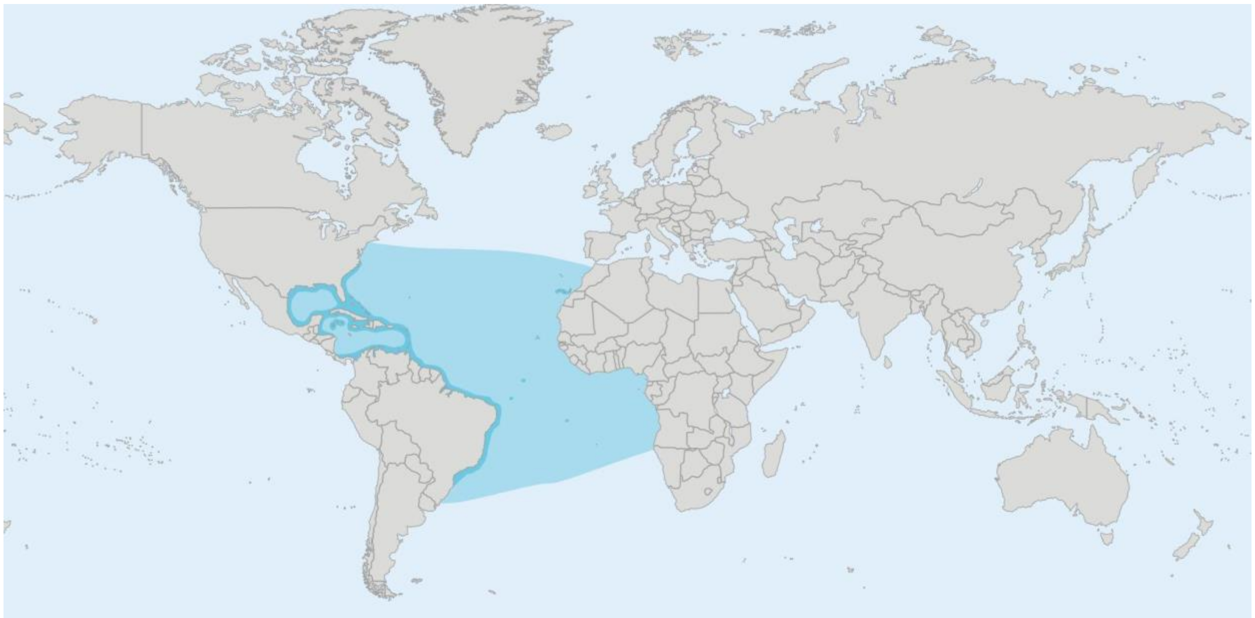

**Figure S1**. Geographic distribution of *Mobula yarae*. Dark-shaded blue areas denote the confirmed species range, and light-shaded blue areas denote the expected species range. Distribution map reproduced with permission from the Manta Trust.

A

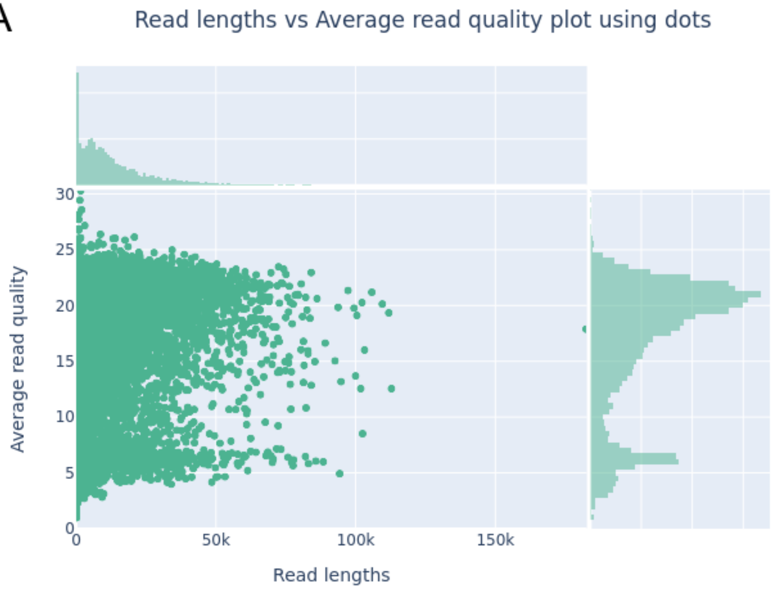


B

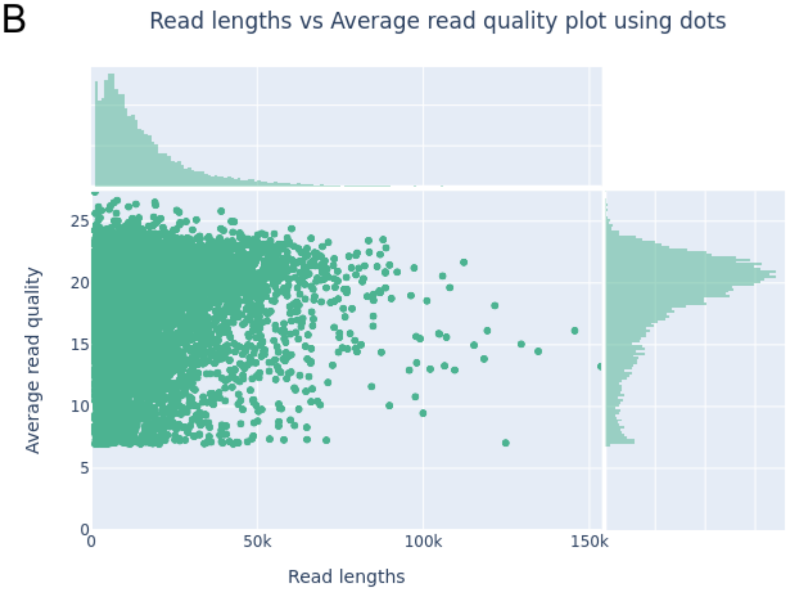


**Figure S2**. Oxford Nanopore Sequencing read quality *for Mobula hypostoma*. Scatterplots of read length versus average read quality for (A) raw and (B) quality filtered sequencing data. Each point represents a single read and is plotted by its length along the x-axis and mean base quality score along the y-axis. Marginal histograms display the distribution of read lengths (top) and average quality scores (right).

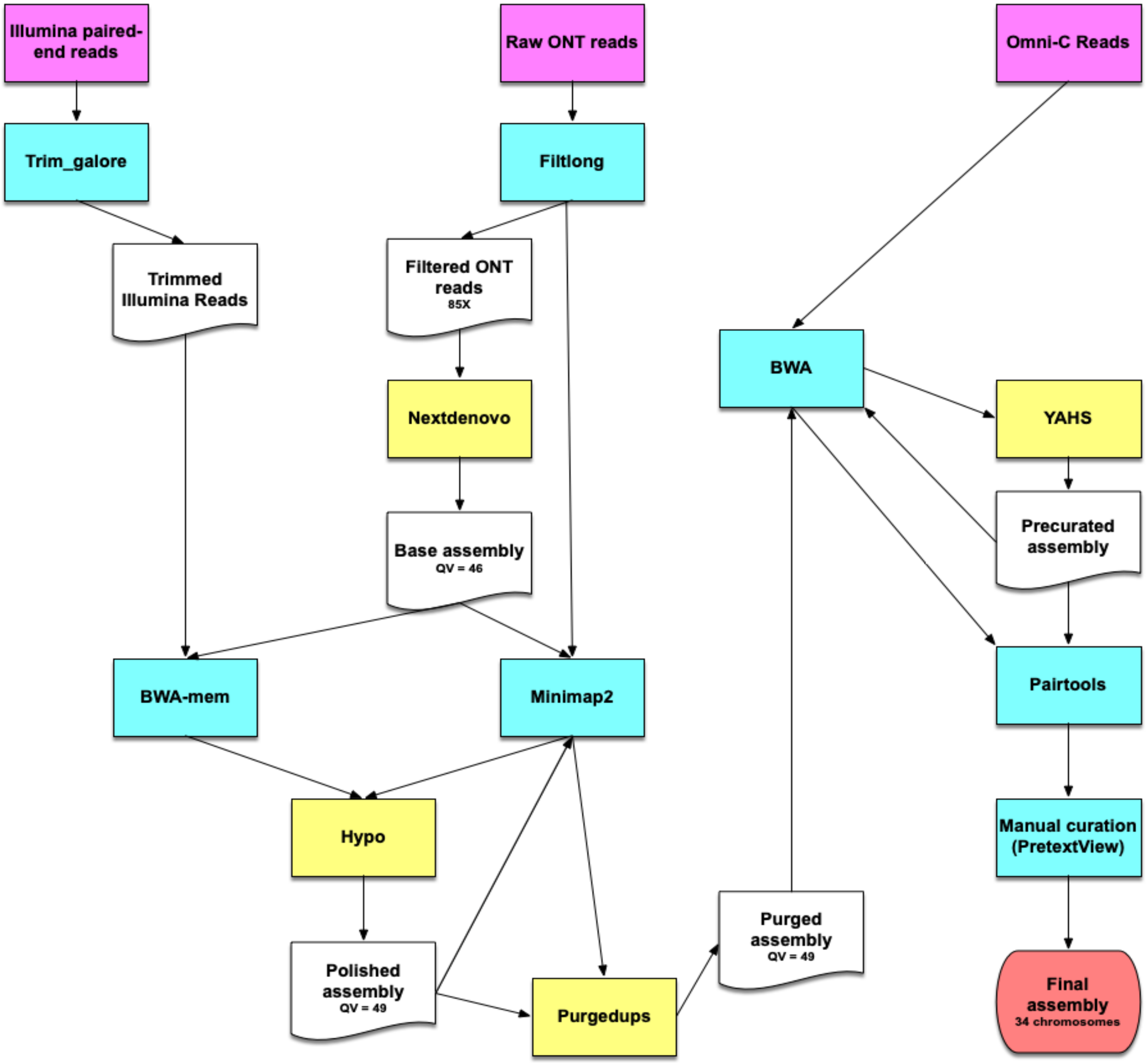


**Figure S3.** CNAG's Long-read Assembly Workflow in Snakemake (CLAWS) used to assemble the *Mobula hypostoma* genome.

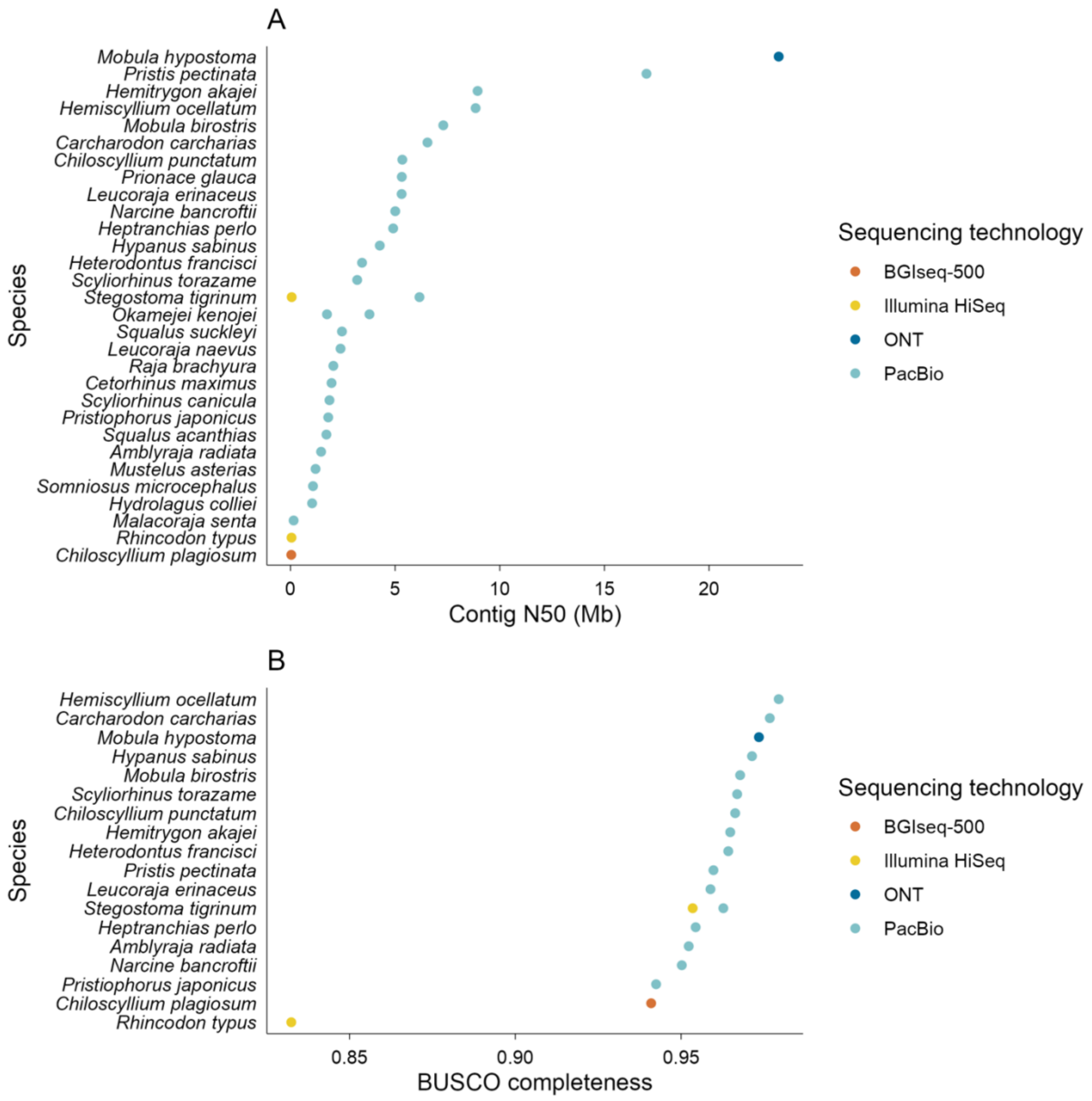


**Figure S4.** Contig N50 (A) and BUSCO completeness scores (B) for the chromosome-level chondrichthyan genome assemblies available on NCBI in October 2025. BUSCO completeness scores are shown for annotated genomes only. Points are coloured according to the sequencing technology used for genome assembly.

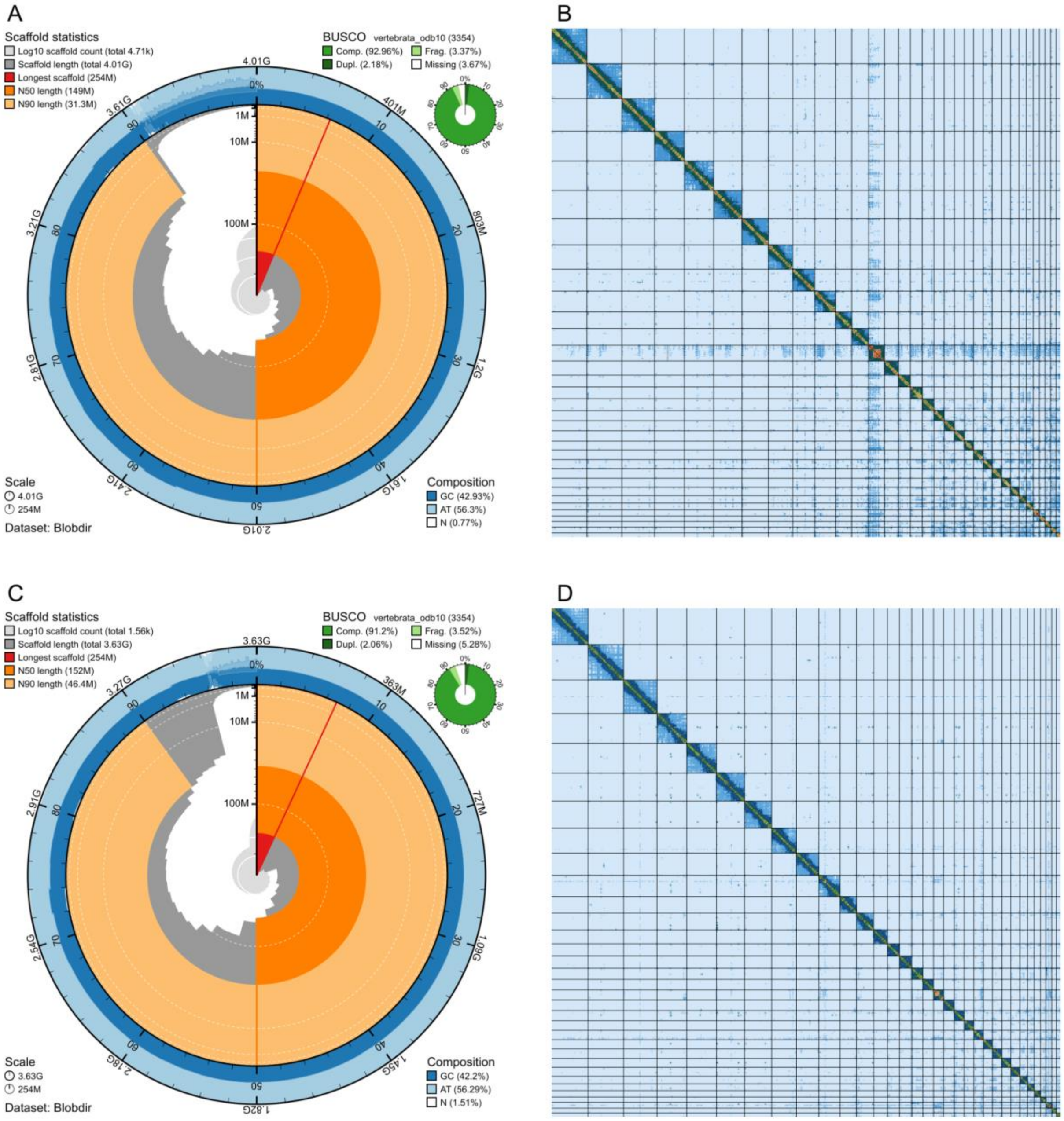


**Figure S5.** Snail plots generated using BlobToolKit to visualize key quality metrics of the *Mobula yarae* haplotype-resolved assemblies sMobBir1.hap1 (A) and sMobBir1.hap2 (C). Tracks include scaffold count, scaffold length, scaffold N50/N90, GC content and BUSCO scores. Hi-C contact maps for sMobBir1.hap1 (B) and sMobBir1.hap2 (D) generated with PretextSnapshot. Chromosomes are ordered and orientated along the axes and the heatmap shows contact frequencies with the strong diagonal signal indicating high contiguity and correct orientation of chromosome-length scaffolds.

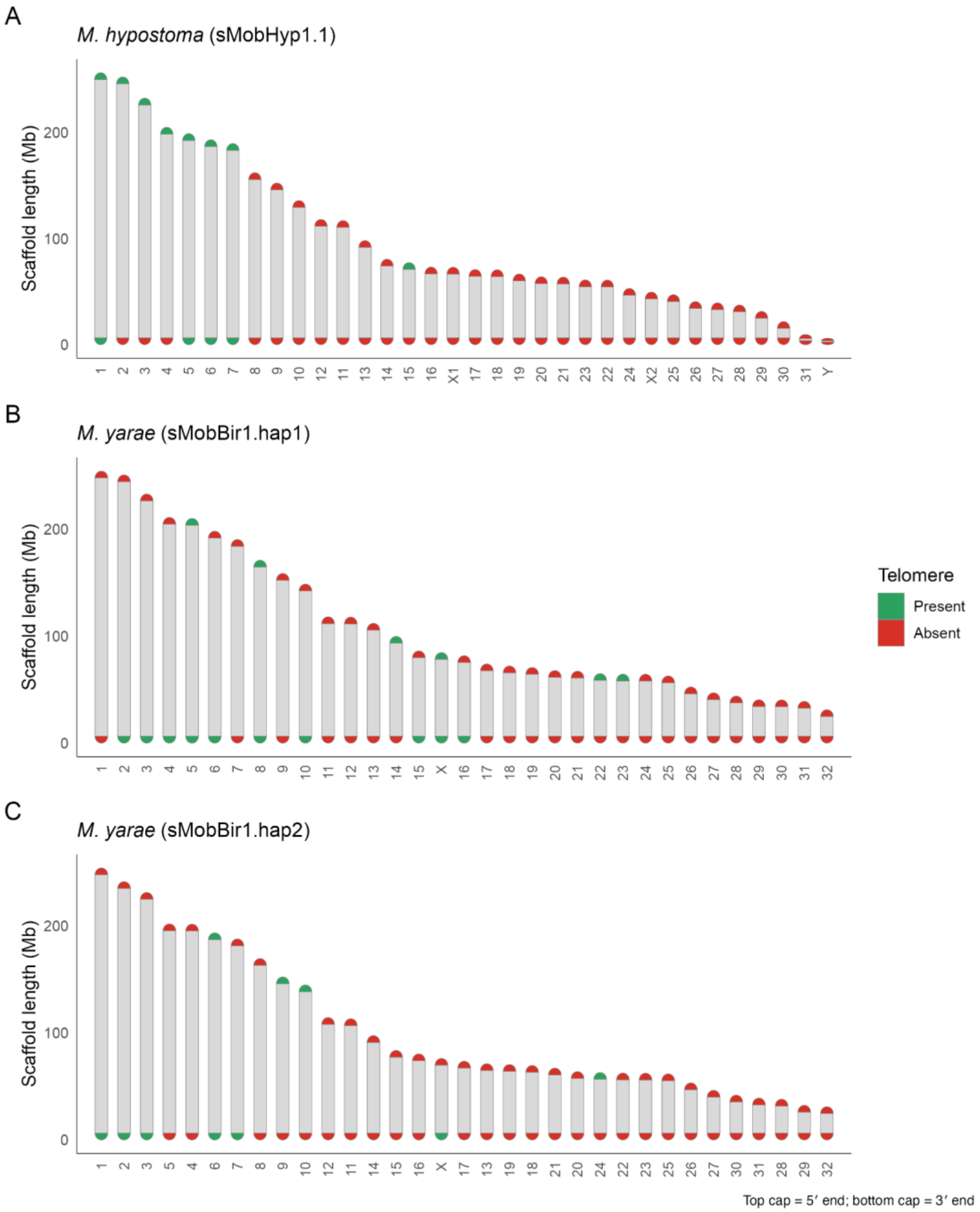


**Figure S6.** Telomere identification in (A) *M. hypostoma*, (B) *M. yarae* haplotype 1, sMobBir1.hap1 and (C) *M. yarae* haplotype 2 sMobBir1.hap2. Each bar represents a chromosome-length scaffold (≥ 5Mb) ordered by length. Telomeres were identified by scanning the terminal 100 kb of each scaffold ≥5 Mb for the canonical vertebrate telomeric motif (TTAGGG/CCCTAA). A scaffold end was classified as telomeric if repeat density exceeded a threshold of 10 per 1,000 bp in any window within the 100 kb terminal region. Upper and lower caps represent the start and end of each chromosome and are shown in green when they passed the telomere detection threshold and in red when they did not.
